# Color and Spatial Frequency Provide Functional Signatures of Retinotopic Visual Areas

**DOI:** 10.1101/2022.12.01.518789

**Authors:** Spencer R. Loggia, Stuart J. Duffield, Kurt Braunlich, Bevil R. Conway

## Abstract

Primate vision relies on retinotopically organized cortex defined by representations of hemifield (upper versus lower visual field), eccentricity (fovea versus periphery), and area (V1, V2, V3, V4). To test for the functional signatures of these organizing principles, we used fMRI to measure responses to colored gratings with varying spatial frequency, color, and saturation across rentinotopically defined parcels in macaque monkeys. Using an approach we call Sparse Supervised Embedding (SSE), we identified stimulus features that differed most across the cortical parcels. Constraining the model according to each voxel’s eccentricity revealed the expected variation of spatial frequency and S-cone modulation with eccentricity. Constraining according to voxel dorsal-ventral location and retinotopic area, revealed two additional components which together provide unexpected functional signatures. First earlier visual areas were more sensitive to contrast and saturation compared to later areas. Second, ventral parcels responded less to luminance contrast and more to color, particularly along the orange-blue intermediate axis in cone-opponent color space. This dorsal-ventral asymmetry resembles differences between candidate dorsal and ventral subdivisions of human V4 but extends to include all retinotopic visual areas, with a magnitude that increased from V1 to V4. Overall, our study demonstrates SSE as a data-driven tool for generating hypotheses about cortical function and behavior, and provides insight into the functional roles of different retinotopic areas.

**Significance Statement:** This study explores the functional roles of visual areas (V1, V2, V3, V4, V3a, MT) parceled by eccentricity preference and visual hemifield. We employ a new approach called Sparse Supervised Embedding (SSE), a promising technique for visualizing and understanding complex neuroimaging datasets. Analyzing fMRI responses to different color grating stimuli, we uncover a functional organization across these parcels, shedding light on their roles in visual computation. This study enhances our knowledge of processing in early visual cortex and how it varies across the retinotopic map. It also aids in resolving apparent differences between prior human and macaque studies.

## Introduction

The distinct computational objectives of the multiple visual areas early in the cortical visual-processing hierarchy (V1, V2, V3, and V4) remain unclear (Zhang et al., 2023). Each area is retinotopically organized (Hubel et al., 1977; Gattass et al., 1988), leaving an imprint on higher-order areas (Hasson et al., 2003; Lafer-Sousa and Conway, 2013; Arcaro et al., 2017; Klink et al., 2021) and influencing various aspects of visual behavior. For instance, acuity is determined by a target’s location relative to the fovea, and perception can vary with the polar angle within the visual field - including salient differences in perception between upper and lower visual field merridians (Carrasco and McElree, 2001; Levine and McAnany, 2005; Quek and Finkbeiner, 2015; Himmelberg et al., 2023). These perceptual asymmetries are not solely explained by retinal factors, suggesting an explation might be found in retinotopic cortex (Kupers et al., 2022).

Yet the distinct roles of different retinotopic areas remains elusive. Specifically, it is unclear whether the multiplicity of lower-order areas serve to build complex receptive fields (Livingstone and Hubel, 1988; Freeman et al., 2013; Vernon et al., 2016; Ponce et al., 2019) or whether they emerge from the goal of smoothly representing the visual field at multiple scales (Tarhan et al., 2021). An understanding of the functional roles of retinotopic areas is further complicated by potential differences between humans and monkeys, particularly concerning V4 (Wade et al., 2002; Hansen et al., 2007; Winawer et al., 2010; Goddard et al., 2011).

To address the different roles that retinotopic areas might play in visual computation, here we ask: to what extent can combinations of simple stimulus features distinguish retinotopically defined parcels? Our objectives are twofold: to advance a quantitative framework for understanding neural responses across the visual-processing hierarchy and their connection to perception (Himmelberg et al., 2023); and to facilitate linking neural activity in monkeys to psychophysical and functional magnetic resonance imaging (fMRI) data in humans (Singh et al., 2000; Wade et al., 2002; Liu and Wandell, 2005; Brewer et al., 2005; Pihlaja et al., 2008; Mullen, 2019).

We first used fMRI retinotopic mapping in macaque monkeys to parcelate visual cortex by eccentricity, upper and lower visual field, and visual area (V1, V2, V3, V4). We then measured fMRI responses to gratings varying in color, spatial frequency, and cone-contrast. We analyze the data with a method we call Sparse Supervised Embedding (SSE) that discovers differences between subdivisions of a dataset in a high-dimensional space. In neuroimaging similar linear methods such as LDA (Linear Discriminant Analysis) are often used to separate stimulus-defined classes by voxel responses (Preul et al., 1996; Cox and Savoy, 2003; Pardo et al., 2006; Lin et al., 2021). We aim to flip this logic, instead uncovering sparse combinations of stimuli that best distinguish retinotopic parcels.

SSE is well suited for this purpose, because rather than simply classifying it finds a space that maximizes variance between the parcels and minimizes variance within them. The axes of this space are linear combinations of features - in this case specific spatial and chromatic gratings - that best separate the cortical parcels. The spatial arrangement of the parcels along these axes is informative about their functional relationships to each other. In our analysis, SSE arranges parcels from central to peripheral along an axis in the stimulus feature space known to vary with eccentricity (Wright and Johnston, 1983; Curcio et al., 1991; Vanston and Crognale, 2018; Broderick et al., 2022). It also produces axes that separate the parcels in other physiologically meaningful ways, including functional asymmetries between the upper and lower visual field.

## Methods

### Functional Magnetic Resonance Imaging Acquisition

Imaging acquisition was the same as in (Lafer-Sousa et al., 2012; Lafer-Sousa and Conway, 2013). Some of the data analyzed presently was used in prior publications, and other data has not previously been analyzed; all the data analyzed in the present report will be available open access at neicommons (https://neicommons.nei.nih.gov). Two alert rhesus macaques (7-8 kg, M1 and M2), were scanned at the Martinos Imaging Center at Massachusetts General Hospital (MGH) in a 3 Tesla Allegra (Siemens, New York, New York) scanner using a custom-made four-channel send-receive surface coil (Martinos Center, MGH, Charlestown, Massachusetts). Images were acquired using standard Echo-Planar Imaging (EPI) methods, with 2 second repetition time (TR), each repetition acquiring a 98 x 63 x 98 voxel matrix with 1mm isotropic voxels. Animals were trained using juice rewards to sit in a sphinx position in a custom-built plastic chair placed inside the bore of a horizontal scanner while fixating a central target on a display screen; animals were required to fixate throughout the experiment to receive reward. Head position was maintained with custom plastic head posts that were surgically implanted (see surgical methods in Lafer-Sousa et al. (2012)). Eye movements were tracked with an infrared eye tracker (ISCAN, Burlington, Massachusetts). Animals were rewarded for maintaining fixation within a degree of the central fixation target. Monocrystalline Iron Oxide Nanoparticle (MION) (AMAG Pharmaceuticals, Cambridge, Massachusetts; 8-10 mg/kg, diluted in approximately equal volume of saline), was injected intravenously in the saphenous vein immediately prior to scanning to improve the magnetic resonance signal (Vanduffel et al., 2001). High-resolution anatomical scans (0.35 mm x 0.35 mm x 0.35 mm in M1 and 0.35 mm x 0.4 mm x 0.35 mm in M2) were obtained while the animals were lightly sedated during a separate scanning session. All imaging and surgical procedures follow local and National Institute of Health guidelines. All imaging and surgical procedures were approved by the Harvard Medical School Institutional Animal Care and Use Committee.

### Experimental Design and Statistical Analysis

Four fMRI experiments were conducted over the course of 7 sessions in M1 and 6 sessions in M2. In experiment 1, we measured responses to vertical and horizontal flickering checkerboard wedges to define retinotopic areas as described in (Lafer-Sousa and Conway, 2013)(Figure 1a). In experiment 2, we measured responses to checkerboard patterns restricted to rings of different eccentricity (Figure 1b). In experiment 3, we measured responses to color-gray gratings in which the colors were defined by a cone-opponent color space (Figure 2a,b) (MacLeod and Boynton, 1979; Derrington et al., 1984); the stimuli varied in hue and saturation (Figure 2c); and in experiment 4, we measured responses to heterochromatic gratings also defined by the cone-opponent color space, but varying in spatial frequency (Figures 2d). Each scan session consisted of 13-24 stimulus runs, details of which are described below.

**Figure 1.**
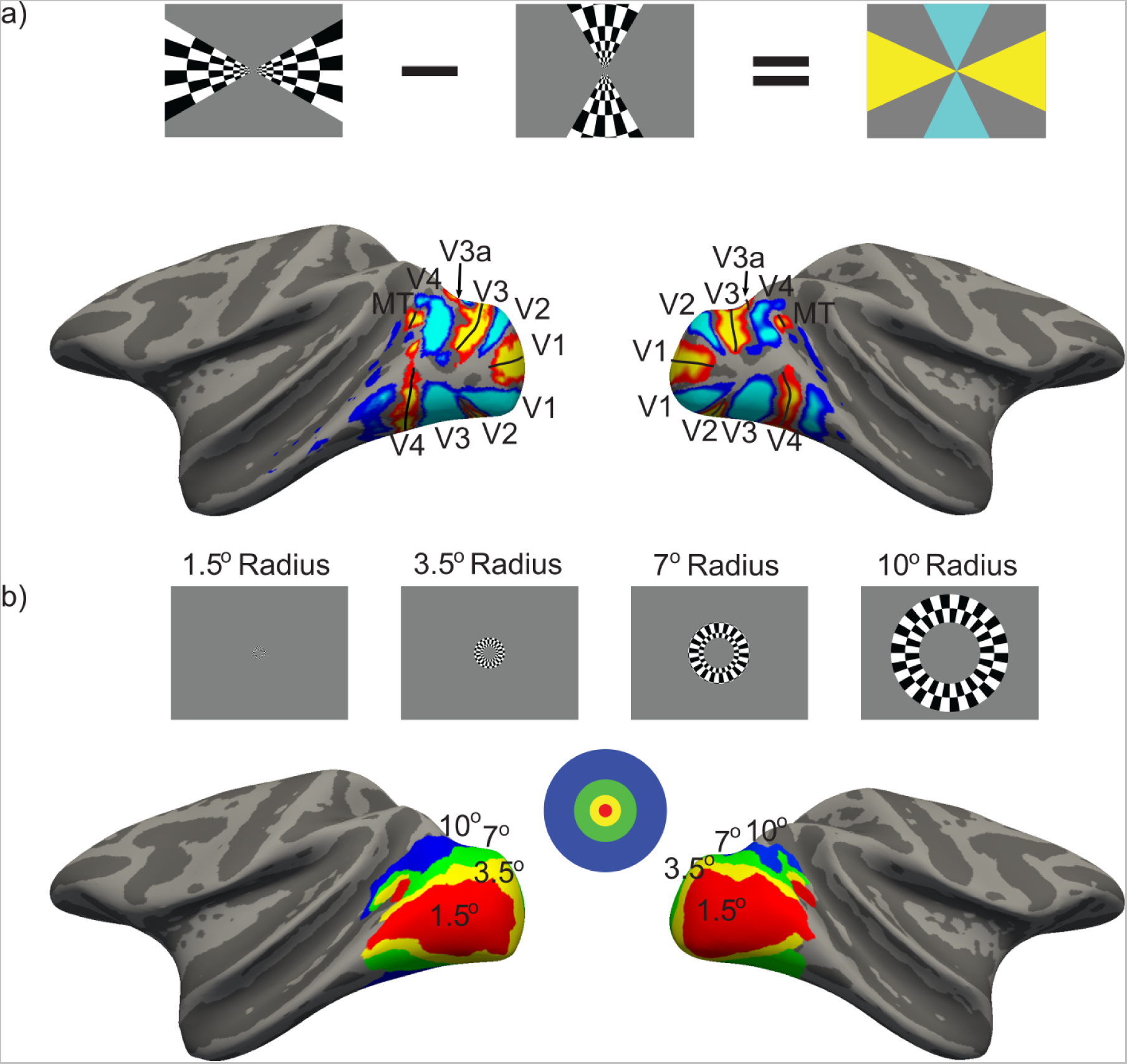
Parcellation of retinotopic visual cortex in macaque monkey using functional data obtained in the same individual animals and functional atlases. a) Individual-specific functional parcellation. To delineate retinotopic areas in each macaque, the animals were shown vertically and horizontally oriented checkerboard wedges in a blocked paradigm (icon). The panel shows the contrast maps between these two conditions displayed on the inflated surface of monkey 1 (M1), which reveals the borders of the retinotopic areas along the vertical (shown in blue) and horizontal (shown in red) meridians; for example, the posteriormost vertical meridian delineates V1 from V2, and the posteriormost horizontal meridian delineates upper from lower V1. b) To define voxels by eccentricity responses, checkerboard rings and circles were presented in a blocked paradigm, and the statistical contrasts between these conditions were used to create parcels encompassing the central 1.5 degrees, and nested annuli extending to 3.5 degrees, 7 degrees, and 10 degrees (icon). The eccentricity parcellation is shown for M1. Note the existence of two representations of the fovea, the larger one corresponds to the confluence of the V1-V4 cluster and the other corresponds to the MT cluster (Kolster et al., 2009).

**Figure 2.**
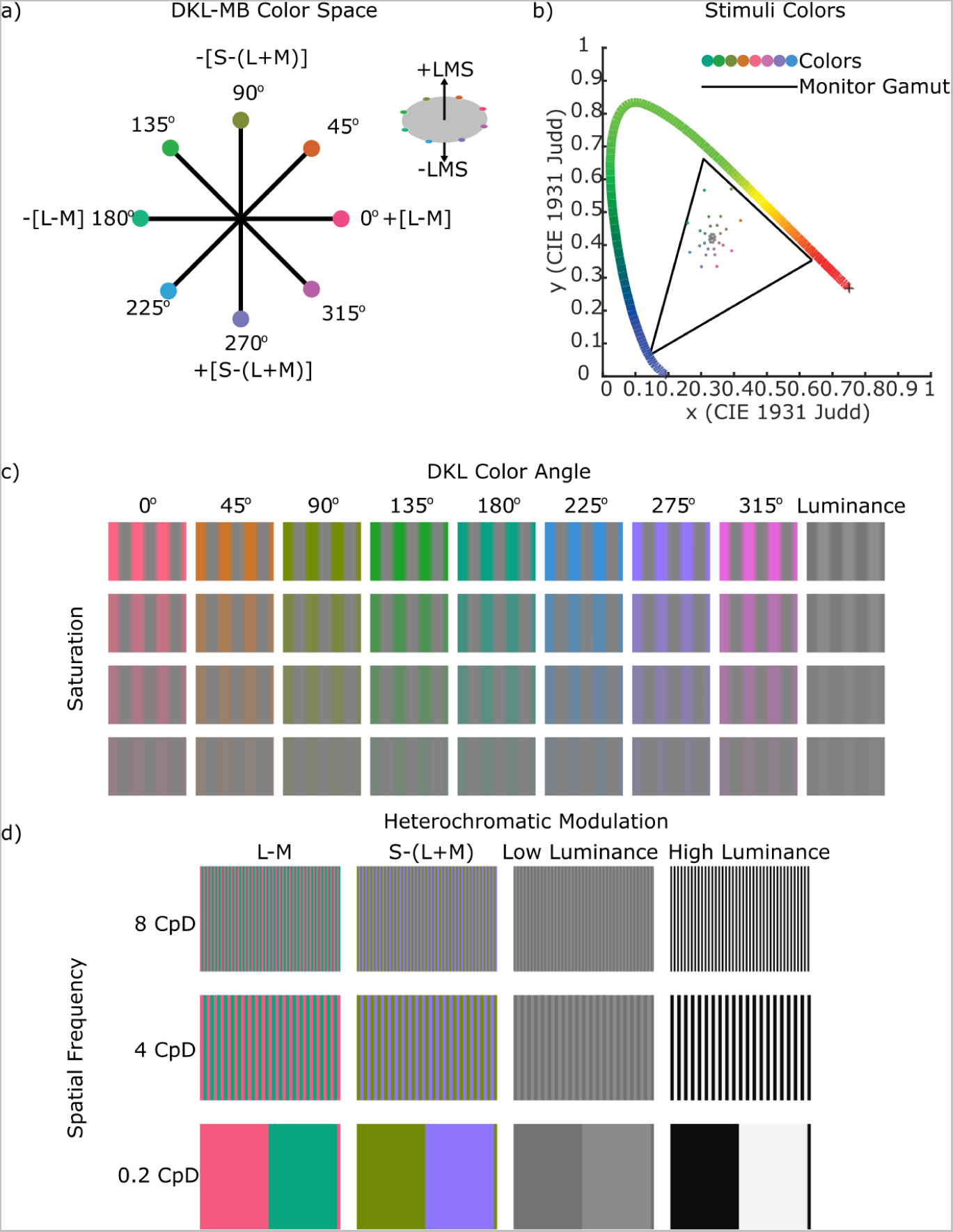
Overview of the stimuli used to identify the functional signatures for retinotopic parcels. a) The colors of the stimuli were defined in a cone-opponent color space. Shown is an equiluminant plane through the space. Colors included those of the two cardinal cone-opponent axes, the two intermediate axes, and the luminance axis (orthogonal to the equiluminant plane) - respectively labelled LM (0 and 180 degrees), Daylight (45 and 225 degrees), S (90 and 270 degrees), Antidaylight (135 and 315 degrees), and Luminance (LMS). b) Plot showing the various saturation levels of the stimuli colors used in one set of experiments (Experiment 3), in the CIE-1931 chromaticity diagram. c) The set of color-gray gratings used in Experiment 3, varying in color angle (columns) and saturation (rows). The spatial frequency of the gratings was 0.5 cycles per degree. d) The set of heterochromatic gratings used in Experiment 4, varying in axis (columns) and spatial frequency (rows, cycles per degree, CpD).

The axes defining the color space consist of two equiluminant cone-opponent axes, one of which is modulated by a change in the difference in excitation values of the L and M cones (the L-M axis, demarcated in Figure 2a by color angles 0 and 180), the other is modulated by the change in excitation value of the S cone (demarcated by color angles 90 and 270), and an orthogonal luminance axis that is modulated by the change in L, M, and S cones (Figure 2a, insert). Colors in each experiment are either modulated along the L-M axis, the S axis, the luminance axis, or two intermediate axes of the equiluminant plane that modulate all three cone types. The colors of the two intermediate axes correspond roughly to the daylight locus (orange/blue; 45 and 225 degrees) and the anti-daylight locus (green/magenta; 135 and 315 degrees). We refer to these five chromatic dimensions as LM, S, LMS, Daylight, and Antidaylight, following others (Goddard et al., 2010; Lafer-Sousa et al., 2012). The gamut of the colorspace was bounded by the monitor gamut (Figure 2b).

The stimuli were presented on a screen 48.2 cm away from the animal, with a JVC DKLA projector (1024×768 pixel resolution). For experiments 1, 3, and 4, the projected image subtended 22.5 cm by 16.9 cm (27 x 20 degrees of visual angle, DvA), and for experiment 2, the projected image subtended 35 cm by 26.8 cm (41 x 31 DvA). The projector’s luminance output was linearized, and colors were measured and calibrated using spectral readings taken from a spectroradiometer (PR-655, Photo Research, Inc, Chatsworth, California). Spectra were multiplied by the Judd-revised CIE 1931 color matching functions to derive CIE xyY coordinates (Figure 2b). All stimuli were presented in a blocked paradigm. For each experiment, each stimulus presentation was matched in duration. Stimulus blocks were interleaved with adapting-gray blocks.

The first experiment (Experiment 1, Figure 1a) allowed us to identify horizontal and vertical meridian representations, which define the boundaries between retinotopic visual areas. Stimuli consisted of black-and-white checkerboard wedges that flickered between complementary checkerboard patterns every 1 second at 99% luminance contrast. The stimuli were two wedges that radiated from the central fixation spot. In one block the wedges spanned the vertical meridian (60 degree wedge); in the other block, the wedges spanned the horizontal meridian (30 degree wedge). Blocks were 32 seconds long (16 TRs) and each run had 16 blocks. For all runs, the conditions were ordered Horizontal, Gray, Vertical, Gray, repeated four times. A total of 13 runs of this experiment were collected in M1, and 14 runs were collected in M2.

The second experiment was an eccentricity mapping experiment (Experiment 2, Figure 1b). Stimuli consisted of LMS, LM, or S checkerboards presented in rings centered on the fixation point; the checkerboards flickered between complementary checkerboard patterns every 1 second. Rings extended from the central fixation spot to 20 DvA, with each successively larger ring having an inner radius equal to the outer radius of the previous ring. Rings had outer radii of 1.5, 3.5, 7, 10 DvA. Blocks showing one of the rings were interleaved with blocks of the adapting gray background. Blocks were 32 seconds (16 TRs) and each run had 17 blocks. There were three different run orders, with only the colors of the checkboards varying across the orders. For all three run orders, the gray blocks and rings proceeded as Gray, 1.5 degrees radius, Gray, 3.5 degrees, Gray, 7 degrees, Gray, 10 degrees, repeated twice, with a final gray block. For the first order, the first four ring blocks were S checkerboards and the second four ring blocks were LM; the second order had S followed by LMS checkerboards; the third order had LMS followed by LM checkerboards. A total of 24 runs were collected in M1 and 21 runs were collected in M2.

In the third experiment, we presented color-gray “trapezoidal” gratings with a spatial frequency of 0.5 cycles per degree (CpD) (Figure 2c). The saturated or fully gray portions of the gratings made up 80% of each cycle; the transition between the fully gray and fully saturated portions of the gratings was progressive such that the cone contrast from one side of the grating to the other had a trapezoidal shape. This spatial structure helps mitigate chromatic aberration. The gratings were presented in nine colors, defined by the eight poles of the four axes in the equiluminant color plane and the luminance axis; each grating was presented at four saturation levels (10%, 30%, 50%, or 95% of the maximum saturation of the display for the colored gratings; 1%, 2.5%, 5.5%, or 8.7% luminance contrast for the achromatic gratings; 36 stimulus blocks total; see Supplementary Table 1 for the cone contrasts for each color). Color pairs along the Daylight axis had the same absolute spatiotemporal Michelson cone contrast as the color-pairs along the Antidaylight axis. The gratings drifted horizontally at a speed of 0.8 cycles per second, and the direction of the drift changed every two seconds. Blocks of gratings were interleaved with blocks of gray. The blocks were 28 seconds (14 TRs) and each run had 19 blocks. Grayscale gratings appeared in the middle of each run. A total of 49 runs were collected in M1, and 58 runs were collected in M2. Each block type was presented between 26 and 28 times.

The fourth experiment presented heterochromatic gratings with a spatial frequency of either 0.2 CpD, 4 CpD, or 8 CpD; the phases of a given grating were defined by the poles of LM axis, the S axis, or the luminance axis (Figure 2d). The colors of the colored gratings were set at 90% of the saturation of the display defined in the DKL-MB colorspace; the achromatic gratings were of either 90% or 9% luminance contrast. Blocks of gratings were interleaved with blocks of gray. During a given block, the grating was stationary and alternated in counter-phase between complementary patterns every one second. The blocks were 32 seconds (16 TRs) and each run had 25 blocks. A total of 14 runs were collected in M1, and 13 runs were collected in M2. Each block type was presented 27 times total.

fMRI data preprocessing is described in the “fMRI Preprocessing” section of the methods. F-tests for linear regressions were computed with MATLAB’s fitlm function. Multi-way ANOVAs, ANCOVAs, and subsequent Tukey’s Honest Significant Difference tests were computed with MATLAB’s anovan, ancova, and multcompare functions. Multi-way ANCOVA was computed using the MANCOVAN matlab toolbox (Gruner, 2010). In cases where bootstrapping was used to approximate confidence intervals, data was independently resampled 1000 times. Variance and expected value were determined by fitting a normal distribution over all resamples. P values and statistical tests used are presented in the figures, figure legends, and results section. The SSE analysis method is described in detail in the “Sparse Supervised Embedding” section of the methods.

### fMRI Preprocessing

The raw data was unpacked from DICOM (Digital Imaging and Communications) to NIfTI (Neuroimaging Informatics Technology Initiative) format using dcm2niix (Li et al., 2016). The images underwent thermal denoising using the NORDIC algorithm (Vizioli et al., 2021) to improve the signal-to-noise ratio. Images were reoriented from the sphinx position. Data was motion corrected with the FSL (fMRI-Brain Software Library) motion correction algorithm (MCFLIRT) with 12 degrees of freedom (Smith et al., 2004). The functional volumes were then co-registered to the anatomical volumes using ITK-SNAP’s linear registration tool v3.6.0 (Yushkevich et al., 2006) and ANTs non-linear registration algorithm (Avants et al., 2011) to get the closest mapping of the functional volumes to the anatomical image. Blocks were one-hot encoded and convolved with the MION hemodynamic response function (HRF) to create design matrices (Vanduffel et al., 2001). No spatial smoothing was applied. Nilearn’s GLM (General Linear Model) module (https://nilearn.github.io/) was used to calculate *β* coefficients and create statistical contrast maps, using drift regressors up to the 3rd order polynomial to account for fMRI signal drift.

### Anatomical Processing and Region of Interest Definition

To create surfaces of the macaque brain, the high-resolution anatomical images were skull stripped, white matter regions were labeled, and surfaces were generated and inflated with FreeSurfer (Smith et al., 2004). Retinotopic parcels were defined using functional data obtained in each animal (Experiment 1); the data are from the two animals used in (Lafer-Sousa et al., 2012; Lafer-Sousa and Conway, 2013). Significance maps showing the contrast of responses to vertical versus horizontal were shown on the inflated surface of each animal; the peak bias of the vertical-greater-than-horizontal responses were used to define the boundary between V1 and V2 and the boundary between V3 and V4; the peak bias of the horizontal-greater-than-vertical responses were used to define the upper and lower visual field representations of V1, and the boundary between V2 and V3. MT+ was defined such that it bordered V4d along a shared horizontal meridian and featured a continuous eccentricity map in a three quarters circle around the foveal representation in the superior temporal sulcus (Figure 1b) (Brewer et al., 2002; Kolster et al., 2009). Surface labels of the retinotopic areas were transformed into the anatomical volume space. These retinotopic areas were cross-referenced with the Paxinos illustrated atlas (Paxinos et al., 2000), the D99 atlas (Reveley et al., 2016), and the CHARM atlas(Jung et al., 2021), producing retinotopic regions of interest (ROI) for each subject (Figure 1). Retinotopic areas V1, V2, V3, and V4 were subdivided into dorsal and ventral ROIs by extending the horizontal meridian of V1 through the center of the foveal confluence on each subject’s surface. These surface labels were then back-projected into the anatomical space and used as a reference to divide the retinotopic ROIs into ventral and dorsal regions, corresponding to the upper- and lower-visual field representations of each retinotopic area. V3a and MT+, being located on the dorsal surface of the cortex, were not subdivided, despite these areas having representations of the upper and lower visual field (Gattass et al., 1988; Kolster et al., 2009; Zhu and Vanduffel, 2019). To determine eccentricity representations, statistical contrasts of the responses elicited by neighboring rings were generated, and voxels were assigned to eccentricities eliciting the maximum z-score, or zero if the z-score was less than 2 (Figure 1e). Together, the parcelation provides the eccentricity, visual area, and upper-versus-lower visual field representation of each voxel.

### Sparse Supervised Embedding Analysis (SSE)

To accomplish this goal, we developed a method that discovers a low dimensional space *E* that best captures the relationships between the retinoptopic parcels, *R*. We do not want the recovered relationships to be unnecessarily constrained, instead we seek a data-driven method that will uncover functional relationships between the parcels. The results should allow us to test the hypothesis that color gratings of varying spatial frequency and contrast provide a functional signature of the parcels. Given a number of voxels *v*, a number of stimulus conditions *m*, and a number component vectors *k*, we find (*m × k*) projection matrix T that takes the input data matrix (in this case a *v × m* standardized *β* coefficient matrix) into a *k*-dimensional euclidean space *E*, such that the pairwise distance between parcel centers is maximized, while minimizing variance within those parcels. The final objective is the sum of the inter-parcels distance term, the negated intra-parcel variance term, and an L1 regularization term over T (equation 1).

This method is similar in some respects to Linear Discriminant Analysis (LDA). LDA finds projection hyperplane that maximizes Fisher’s criterion, i.e. the ratio of inter-class variance to intra-class variance analytically (Xanthopoulos et al., 2013). Our objective function is related, but differs in three key ways. First, LDA is primarily a classification method, and projects *c* classes of input data into a space with *c −* 1 orthogonal dimensions. Equivalently, this can be thought of as finding *c −* 1 hyperplanes that best separate the classes. Since we solve our objective function numerically, and care about the relationships between classes rather than being able to classify them directly, we specify the dimensionality of the embedding space. Second, to encourage sparsity, we add L1 regularization of discriminant vectors. Third, instead of between-class variance, we maximize the pairwise euclidean distance between each class centroid. This allows for control over which class pairs are separated by defining a weighting parameter over the pairwise distance matrix.

After optimization, SSE gives us two useful pieces of information. First, the relative locations of the parcels in *E*. Any reliable structure in the arrangement of the parcels is of interest, because the objective function itself makes no attempt to enforce any relationship between them besides being as statistically different from each other as possible. Second, the weights given to the input features. These indicate the features most useful for statistically separating the parcels. These two pieces of information together provide information about the global relationships between all the parcels with respect to the stimulus features. Specifically, we are interested in the extent to which responses to simple color gratings are informative about the organization of visual cortex by eccentricity, upper v.s. lower visual field, and retinotopic area.

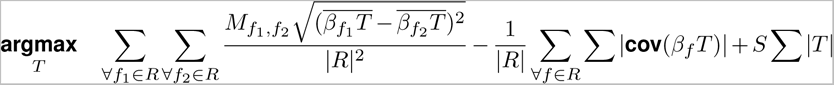

In equation 1, *R* is the set of parcel labels (and *|R|* is the number of parcels), *T* is the transform we are trying to discover (the set of *k* length m component vectors), *β* is the beta matrix with *m* features, *k* is the number of target components or dimensions, and *M* is a weight matrix that allows control over the importance of separating each pair of classes. *f*_1_ and *f*_2_ are used to index specific parcels. We use **cov** to denote computing the covariance matrix over the embedding dimensions. *S* scales the magnitude of the L1 regularization term, used to reduce overfitting, reduce variability between independent runs, and encourage more interpretable component vectors. For all our analyses, *S* is set to 0.005. The model is optimized via gradient descent, using the ADAM optimization method (Kingma and Ba, 2017) as implemented in PyTorch v.1.13. Estimates of error in the SSE models were established by bootstrapping.

*β* coefficients of the stimuli from experiments 3 and 4 in both monkeys were standardized and concatenated into a Voxels × Conditions (*v × m*) matrix. Each voxel was assigned to a retinotopic area and an eccentricity preference. The SSE model was fit on the left and right hemisphere data independently for each animal and across all eccentricities (all indices of matrix *M* are 1).

We ran two further SSE analyses that differ in *M*. First, we maximized the separability of different ROIs across the four eccentricity parcels. To do this, we set *M* such as to not consider differences across the different retinotopic areas either within or between eccentricity levels; the only relationships considered were between different eccentricity levels of the same retinotopic area. This allowed us to find one simple component that best discriminates between more central and more peripheral parcels.

Second, we set *M* to ignore differences across eccentricity, only considering the pairwise distance between all retinotopic areas within each eccentricity level. This has the effect of limiting differences between ROIs due to variation in the size of eccentricity representations.

The top two most positive and top two most negative features for components 1 and 2 were contrasted and projected onto the inflated surface of M1, in order to show the topography of the response to the different component vectors. In these contrasts, the contributions of the components were equal and did not reflect the weighting of each feature in the SSE model.

Table 1 lists the SSE conditions and their corresponding stimuli (see Figure 2). The lowest saturation stimuli, lowest luminance stimuli, and the stimuli of intermediate spatial frequency were excluded from being incorporated as SSE conditions because they were not expected to be informative.

**Table 1.** Conditions corresponding to feature names used in display of SSE discriminant / component vectors.

| Feature Nomenclature | Grating Description(s) |
| --- | --- |
| LM_high_sat | DKL 0 95% Saturation, DKL 180 95% Saturation |
| LM_med_sat | DKL 0 50% Saturation, DKL 180 50% Saturation |
| LM_low_sat | DKL 0 30% Saturation, DKL 180 30% Saturation |
| S_high_sat | DKL 90 95% Saturation, DKL 270 95% Saturation |
| S_med_sat | DKL 90 50% Saturation, DKL 270 50% Saturation |
| S_low_sat | DKL 90 30% Saturation, DKL 270 30% Saturation |
| Daylight_high_sat | DKL 45 95% Saturation, DKL 225 95% Saturation |
| Daylight_med_sat | DKL 45 50% Saturation, DKL 225 50% Saturation |
| Daylight_low_sat | DKL 45 30% Saturation, DKL 225 30% Saturation |
| Antidaylight_high_sat | DKL 135 95% Saturation, DKL 315 95% Saturation |
| Antidaylight_med_sat | DKL 135 50% Saturation, DKL 315 50% Saturation |
| Antidaylight_low_sat | DKL 135 30% Saturation, DKL 315 30% Saturation |
| LMS_high_sat | LMS 8.7% Luminance Contrast |
| LMS_med_sat | LMS 5.5% Luminance Contrast |
| LMS_low_sat | LMS 2.5% Luminance Contrast |
| LM_high_freq | LM 8 CpD |
| LM_low_freq | LM 0.2 CpD |
| S_high_freq | S 8 CpD |
| S_low_freq | S 0.2 CpD |
| LMS_high_freq | High Luminance 8 CpD |
| LMS_low_freq | High Luminance 0.2 CpD |

### Univariate Analysis

For univariate analyses, *β* coefficients for each condition were calculated for each run in which that stimulus condition was presented. Statistical tests used the *β* coefficients from each run as individual observations. Percent signal change for each condition was calculated by dividing the *β* coefficient of the condition by the *β* coefficient of the baseline and multiplying by 100.

We computed features that describe the extent to which each voxel was modulated by the LM, S, Daylight, and Antidaylight color conditions using the fMRI responses to experiment 3. Each condition was estimated by averaging the *β* coefficients computed from the two color-gray gratings that comprise the feature. For example, the LM feature was calculated using responses elicited by the 0-degree (pink-gray) grating and the response elicited by the 180-degree (cyan-gray) grating. Pairs of gratings that made up these features always appeared within the same run at the same saturation, responses to them were averaged for subsequent statistical analysis.

The contrast response function slopes and errors shown in Figure 7a were estimated by fitting a linear regression model to the signal change of each feature as a function of saturation for each ROI, repeated for each bootstrapped sample. The statistical tests comparing the slopes were computed using a one-way ANCOVA. For Figure 7c, the difference in luminance bias for dorsal versus ventral parcels was computed by subtracting the bootstrapped luminance response from the bootstrapped color response for both ventral and dorsal regions, and then subtracting the dorsal difference from the ventral difference.

### Surgical Procedures

Surgical methods are the same as described in (Lafer-Sousa et al., 2012). All surgical procedures follow local and National Institute of Health guidelines. All surgical procedures were approved by the Harvard Medical School Institutional Animal Care and Use Committee.

## Results

To what extent can retinotopic parcels be distinguished by fMRI responses to simple grating stimuli that vary in spatial frequency, color, and contrast? To answer the question, we fit a Sparse Supervised Embedding (SSE) model to responses to these stimuli parcelled by retinotopic region, upper v.s. lower visual field, and eccentricity preference of each voxel. The SSE model finds a sparse linear transformation (*T*) of the *β* coefficients for each stimulus condition that maximizes the pairwise distance between the mean of each parcel while minimizing variances within each class (see methods). As described in the methods, the outcome of the SSE model discovers a low dimensional stimulus feature space that maximally separates the retinoptopic parcels, the resulting organization of the parcels is informative about their functional relationships.

### Functional signatures of eccentricity responses

We first used the SSE model to determine a functional signature of eccentricity, irrespective of the retinotopic area. To do this we disregarded the separation of visual area ROIs - setting the indexes in *M* that correspond to pairwise distances between ROIs within the same eccentricity level to zero. The result yields a single component along which the eccentricity parcels are systematically aranged. The feature contributions to the discriminating component (Figure 3b) show that more foveal voxels were defined by higher weighting for responses to high spatial frequency (LM) gratings and high saturation Antidaylight gratings, while more peripheral voxels were characterized by higher weighting for responses to low spatial frequency (LMS) gratings and high saturation S gratings. These results are consistent with the expectation that foveal representations are tuned for higher spatial frequencies compared to peripheral representations in visual cortex of both humans (Henriksson et al., 2009; Broderick et al., 2022) and monkeys (Schiller et al., 1976; De Valois et al., 1982). Peripheral representations, meanwhile, will be more responsive to S-cone signals given that the relative proportion of S-cones (Curcio et al., 1991) and the apparent saturation of S-cone-modulating stimuli (Vanston and Crognale, 2018) increases with eccentricity.

**Figure 3.**
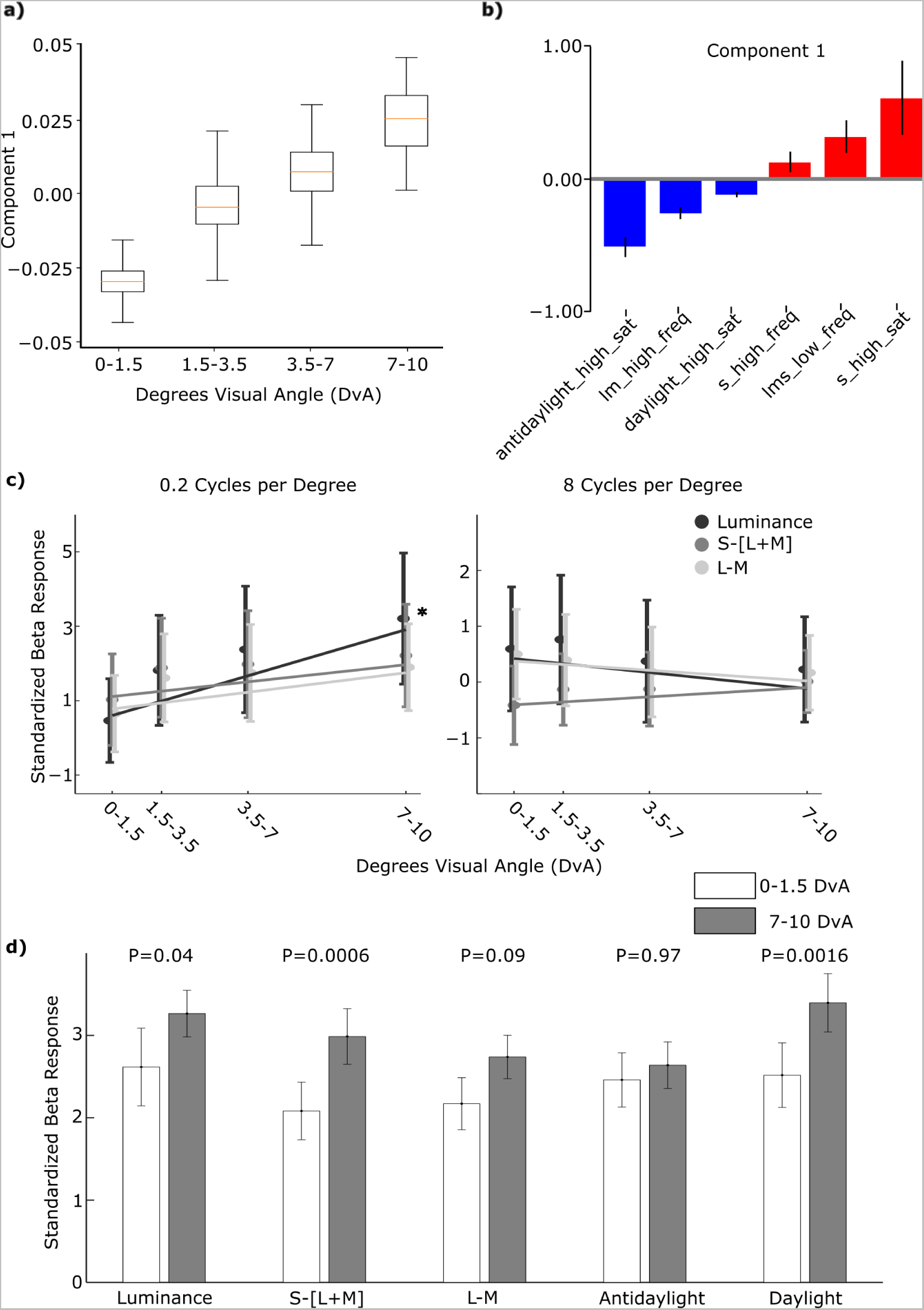
Functional signature of cortical representations of eccentricity recovered with SSE using responses to colored gratings varying in hue, saturation, and spatial frequency. a) One SSE model was fit on all fMRI data (Experiments 3 and 4) obtained in two macaque monkeys. Masking parameter M was set to constrain the model to separate parcels by eccentricity preferences of the voxels, and not their retinotopic area or hemifield (upper/lower) representation (see methods). The response to the component increases with eccentricity (linear regression slope=.02, p=8.4e-16) b) The feature weights for the eccentricity-discriminating component. Features given weight zero are not shown. High saturation S gratings, low frequency Luminance gratings, and high frequency S gratings have strong positive weights (and correspond to more peripheral voxels), while high saturation Antidaylight gratings, high frequency LM gratings, and high saturation daylight gratings have strong negative weights (and correspond to more central voxels). c) The response to heterochromatic gratings varying in color (Luminance, LM, and S) and spatial frequency (0.2 cycles per degree and 8 cycles per degree), across different eccentricity-preferring parcels. Linear regression of the response functions:For 0.2 CpD, Lum slope=0.20, p=0.03 S-[L+M] slope=0.07, p=0.3 L-M slope=0.08, p=0.2; for 8 CpD, Lum slope=−0.5, p=0.5;, S-[L+M] slope=0.03, p=0.5 L-M slope=-0.03, p=0.5; F-tests, DoF=106. d) The response to the highest saturation color-gray gratings (0.5 CpD) within the most foveal (0-1.5 DvA) and the most peripheral (7-10 DvA) cortical parcels. Luminance, S-[L+M], L-M, and Daylight responses increase from fovea to periphery, while Antidaylight does not (Periphery-Foveal Response: Luminance=0.8, p=0.004; S-[L+M]=1.1, p=0.0006; L-M=0.7, p=0.09; Antidaylight=0.3, p-value=1; Daylight=1.01; p=0.002; Tukey’s HSD).

The results of the SSE model are multivariate; to compare the results with those of published reports that use univariate methods, we computed a measure of the responses to the various stimulus conditions as a function of eccentricity. Figure 3c shows how responses to the 0.2 CpD and 8 CpD gratings vary as a function of eccentricity, for the three types of gratings defined by the three cardinal directions in cone-opponent color space (luminance, S, and L-M). Responses to low frequency luminance gratings increased from fovea to periphery (slope=0.19, p=0.03, F-test (DoF=106))—this result is consistent with an increase in response to low spatial frequency luminance gratings from fovea to periphery observed in human visual cortex (Bayram et al., 2016). The responses to the other cardinal directions showed a similar trend, albeit insignificant (For 0.2 CpD, S slope=0.07, p=0.34, L-M slope=0.08, p=0.21; for 8 CpD, Lum slope=-0.045, p=0.46, S slope=0.026, p=0.47, L-M slope=-0.03, p=0.49; F-tests (DoF=106)). For comparison, results obtained in humans with fMRI show gradual increases in responses for low-frequency (0.27 CpD) chromatic gratings as one goes from the fovea to the periphery and gradual decreases in responses to higher-frequency (4.4 CpD) L-M, but not S, gratings (D’Souza et al., 2016).

To begin to relate those perceptual measurements to cortical responses, we investigated the univariate responses across eccentricity to the highest saturation low spatial frequency gratings. In Figure 3d we plot the responses to the highest-saturation color-gray gratings (0.5 CpD) for foveal-preferring voxels (0-1.5 DvA) and peripheral-preferring voxels (7-10 DvA). Response to Luminance, Daylight, and S gratings were higher for the fovea-preferring voxels, while responses to anti-daylight and LM gratings were not different for fovea-preferring and periphery-preferring voxels (t-test, Response of foveal-vs-peripheral voxels: Luminance p=0.004;; S-[L+M], p=0.0006; L-M, p=0.087; Antidaylight, p=0.97; Daylight, p=0.0016; Tukey’s HSD). For comparison, (Vanston and Crognale, 2018) found that suprathreshold stimuli in the periphery, compared to the fovea, were perceived with lower contrast for LM stimuli and higher contrast for S stimuli. The results in Figure 3d are not inconsistent with those findings, but suggest that the differences in color perception between fovea and periphery may be better explained by asymmetries in perception of the intermediate directions in color space rather than asymmetries in perception of the cardinal directions: responses to colors modulated along the daylight axis were different for fovea-preferring voxels versus periphery-preferring voxels, while colors modulated along the antidaylight axis were not. Note that stimuli defined by the two intermediate axes have the same absolute L-M and S cone contrast, so the difference in the responses for stimuli defined by the two intermediate axes through color space cannot be explained by variation in S cone density or macular pigmentation across the retina.

The univariate analyses and SSE uncover patterns of response largely expected from fMRI in humans. Responses in V1 elicited by LMS, S-[L+M], and L-M gratings with a spatial frequency of 0.55 CpD increases as one goes from the central 2 degrees to 8-10 DvA (D’Souza et al., 2016). Luminance gratings cause the greatest increase in response from foveal-preferring voxels compared to peripheral-preferring voxels, while S-[L+M] gratings elicit the smallest increase. Mullen et al., 2007 show that S-[L+M] gratings, also with a spatial frequency of 0.5 CpD, elicit the greatest increase in response from 1 DvA to 6 DvA-preferring voxels of human V1. They also report a decrease in response elicited by L-M gratings across eccentricity.

As far as we are aware, no data has been published comparing responses to Antidaylight and Daylight gratings across eccentricity in human subjects. Goddard et al. (2010) did find generally higher fMRI responses to Antidaylight gratings than Daylight gratings across the visual cortex. The stimuli in that study were somewhat different from those used presently. They were presented on a luminance-contrast pedestal (not equiluminant), were lower mean luminance, increased in spatial frequency from the center to the edge, and were circular. We hypothesize that, controlling for methodological differences, human visual cortex would show an increase in response from foveal to peripheral-responsive voxels for Daylight gratings but not Antidaylight gratings. But univariate methods might not be the most sensitive method for evaluating hypotheses about response variation across eccentricity. The multivariate SSE differences in responses of visual cortex across eccentricity were more substantial than differences in univariate analyses (e.g., compare Figure 3a with Figure 3c).

### Functional signatures of retinotopic visual areas

We next used the SSE approach to test for functional signatures of the retinotopic visual areas and their dorsal versus ventral subdivisions. To do this, we fit the SSE model with two components using only information within voxels matched for each eccentricity preference (Figure 4), which removes the dominant impact of eccentricity on the voxel responses. The results of this analysis recovers a striking pattern. The first component separated parcels along the putative visual-processing hierarchy, from left-to-right in the plot: V1, V2, V3, V4 (upper triangle, circle, inverted triangle, cross). The first component grouped the two dorsal areas, V3a (diamond) and MT+ (square), with the dorsal subdivisions of the other retinotopic areas. The second component separated the ventral and dorsal subdivisions of each visual area: V1v, V2v, V3v, and V4v showed more negative weight compared to V1d, V2d, V3d, and V4d. Areas V3a and MT+ showed more positive values along component 2, again consistent with their dorsal location. The difference between ventral and dorsal parcels along component 2 was progressively greater along the visual-processing hierarchy (Figure 4e) (slope=0.12, p=2.3e-18).

**Figure 4.**
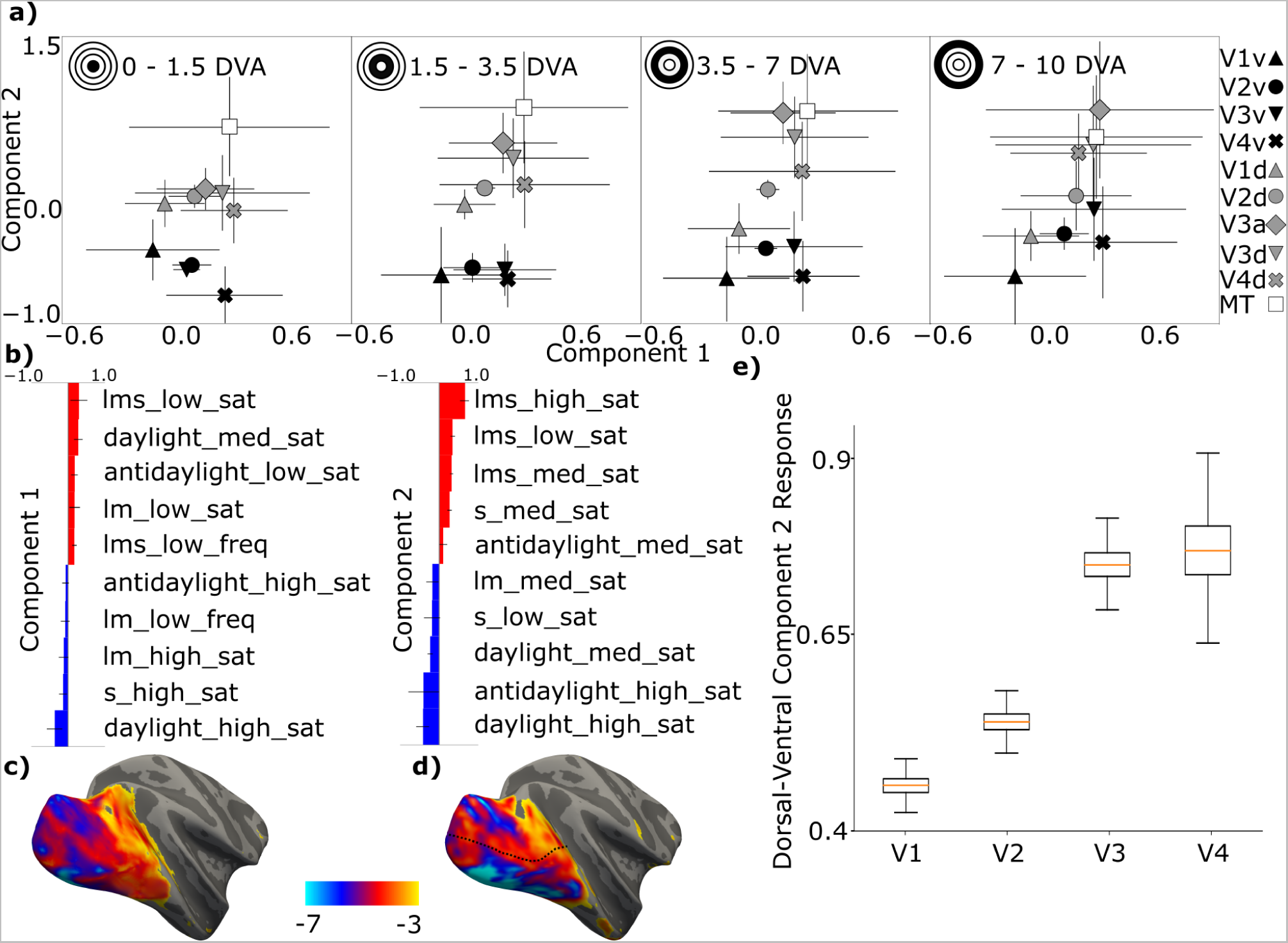
Functional signature of cortical representations of retinotopic areas and their dorsal/ventral subdivisions recovered with SSE using responses to colored gratings varying in hue, saturation, and spatial frequency. a) The mean voxel response for retinotopic areas and their dorsal/ventral subdivisions along the two maximally discriminating components discovered by the SSE model, for voxels defined by eccentricity preference (most foveal-preferring in the leftmost panel; most peripheral-preferring in the rightmost panel. Error bars are 95% confidence intervals via 1000 bootstrapping iterations). The model separates visual area parcels from posterior to anterior along component 1, and from ventral to dorsal along component 2. b) The feature contributions of Components 1 and 2, the two maximally discriminating components. Features given weight zero are not shown. Component 1 positively weights responses to low and medium saturation gratings and negatively weights responses to high saturation gratings. Component 2 positively weights responses to Luminance modulating gratings and negatively weights responses to Daylight, and less significantly Antidaylight, modulating gratings. c) A surface projection of the effect-size contrast between the two features in Component 1 with the most positive weights (Daylight medium saturation, Luminance low saturation) and the most negative weights (Daylight high saturation, S high saturation). d) A surface projection of an effect-size contrast between the two features in Component 2 with the most positive weights (Luminance high saturation, Luminance medium saturation) and most negative weights (Daylight high saturation, Antidaylight high saturation). e) The difference in the Component 2 response between dorsal and ventral subdivisions of each retinotopic area. Whiskers represent 95% confidence intervals estimated via 1000 bootstrapping iterations. The difference increases along the visual hierarchy (linear regression, slope=0.12, p=2.3e-18)

Figure 4b shows the feature contributions for the two components. Component 1 predominantly reflects differences in response to high versus low saturation stimuli. Component 2 predominantly reflects differences in responses to color versus luminance, particularly responses to Daylight-axis colors. To visualize the results, we projected on the cortical surface of the right hemisphere of M1 a contrast between the two most positive and two most negative features of components 1 (separating visual area, Figure 4c), and a contrast between the two most positive and two most negative features of components 2 (separating dorsal and ventral subdivisions, Figure 4d;. the dotted line is the boundary between the dorsal and ventral parcels).

We then used univariate measures to explore the SSE results of the functional signatures of visual areas and their dorsal/ventral subdivisions. Since the component that separates anterior from posterior parcels depends on strong modulation by saturation (negative weights for high cone-contrast gratings and positive weights for low-cone contrast gratings), we analyzed the fMRI responses to the different saturation stimuli as a function of visual area and dorsal/ventral subdivision (Figure 5). We determined the slope of the contrast response functions (CRFs) for stimuli defined by each color axis within each visual area (slopes were a line fit to the fMRI response as a function of contrast, see Methods; Figure 5a). There was a main effect of parcel on the CRF slope; CRFs for all color axes were progressively shallower from posterior to anterior visual areas, as expected by the multivariate results of the SSE.

**Figure 5.**
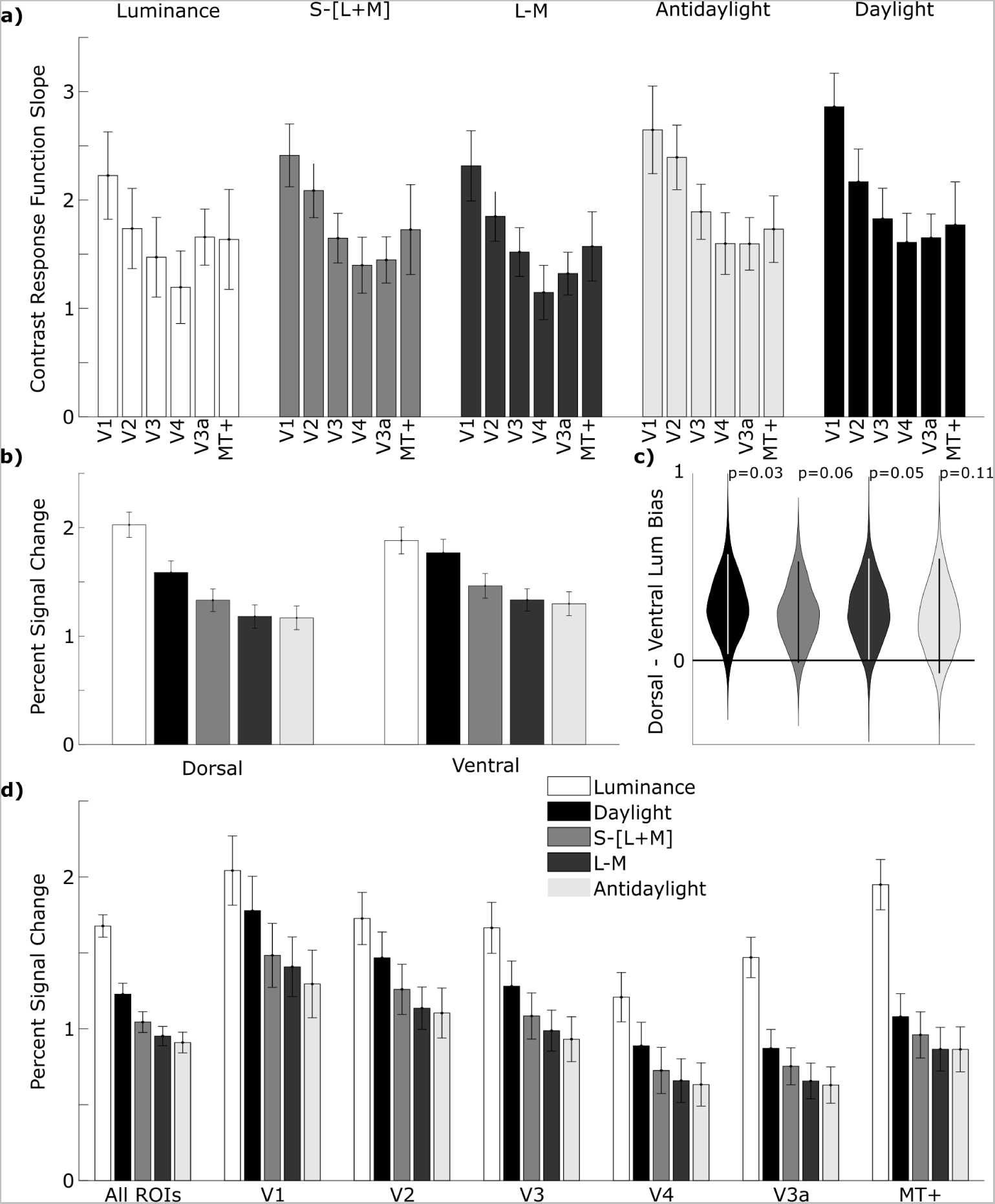
Univariate analyses of the responses of retinotopically defined parcels to colored gratings varying in saturation. a) Contrast response function slope for the fMRI responses to colors defined by various directions through color space (top labels) decrease from V1 to V4 (error bars represent 95% C.I.); CRFs for V3a and MT are shown on the right. The CRF is progressively shallower for areas from posterior to anterior. b) The percent fMRI signal change in response to the gratings defined by various directions through color space (key) for the dorsal and ventral subdivisions of V1, V2, V3, and V4. Each bar shows the average response to the three gratings of highest saturation for each color axis (see Figure 3c). The responses to gratings modulated along the Luminance axis were significantly greater than the responses to gratings modulated along the Daylight axis in dorsal parcels, but not ventral parcels. (2-way ANOVA, main effects of color, F=58, p=0; region, F=6.3, p=0.01; and an interaction of region and color, F=2.7, p=0.03; post-hoc test comparing response to Luminance gratings versus Daylight gratings, Tukey’s HSD, Dorsal Luminance - Dorsal Daylight: p=2.8e10-6, Ventral Luminance - Ventral Daylight: p=0.93) c) Violin plot comparing dorsal versus ventral regions for the difference between the responses to gratings modulating along the Luminance axis versus the responses to gratings modulating along the other color axes (from left to right: Daylight, S, L-M, and Antidaylight). Dorsal parcels show a bias for Luminance regardless of what color is used as the comparative response (note the p values in the plot), but the bias is greatest when compared with responses along the Daylight axis. d) The percent signal change for gratings defined by each direction through color space, combined across all saturation levels. Gratings along the Daylight axis elicited a greater response than colored gratings along all other axes (excluding luminance) for responses averaged across all retinotopic parcels (left plot: Daylight versus S, p=0.001; Daylight versus LM, p=8.9e-10; Daylight versus Antidaylight, p=2.9e-10). Right panels show this analysis for each visual area separately (multiple comparisons render statistical tests insignificant).

The SSE model also suggests that differential responses to Daylight and Luminance gratings separate dorsal and ventral parcels. To evaluate this directly, we plot the fMRI responses to each color grating averaged across the 4 saturation levels for dorsal vs. ventral parcels (Figure 5b). There is a difference in fMRI response in the dorsal and ventral parcels that is modulated by color (2-way ANOVA, main effects of color (F=58, p=0), region (F=6.3, p=0.01), and to a lesser extent interaction of parcel and color (F=2.7, p=0.03)). Post-hoc tests reveal that although individual color responses do not differ between dorsal and ventral parcels (Tukey’s HSD, p-values > 0.05), responses to Luminance-contrast gratings were greater than responses to Daylight gratings for dorsal but not ventral parcels (Tukey’s HSD, Dorsal Luminance - Dorsal Daylight: p=2.8e-6, Ventral Luminance - Ventral Daylight: p=0.93).

To investigate the increase in color response relative to luminance response in the ventral subdivisions compared to the dorsal subdivisions, we computed a univariate feature guided by the SSE results. We determined the difference in luminance bias between ventral and dorsal parcels (Figure 5c), where the luminance bias was computed as the difference in response to luminance gratings and the response to each of the different colored gratings. The Dorsal parcels showed a luminance bias regardless of which color grating was used to assess it, although not all of the comparisons were significant (Daylight-versus-Luminance, p=0.03; LM-versus-Luminance, p=0.05; S-versus-Luminance, p=0.06; Antidaylight-versus-Luminance, p=0.11; one-sided bootstrap t-tests).

The signatures of retinotopic areas and their dorsal/ventral subdivisions recovered by SSE consistently assigned high weight to the responses to Daylight-axis modulating gratings (three of the four highest weight features in components 1 and 2 implicated gratings that modulated along the Daylight axis, Figure 4). So, we investigated whether cortical responses were higher to gratings that modulated along the Daylight axis compared to gratings that modulated along other directions in color space. Across all retinotopic parcels, the average fMRI response was highest to luminance-modulating gratings; Among the responses to colored gratings, the highest response was to gratings that modulate along the Daylight axis (Figure 5d left panel, Daylight versus S, p=0.001; Daylight versus LM, p=9×10-10; Daylight versus Antidaylight, p=3e-10). This pattern was consistent in all visual areas (Figure 5d, right panels), but varied somewhat across areas (2-Way ANOVA, main effects of both area and color axis; F=85, p=0.0 for ROI, F=61, p=0.8 for Color Axis). Figure 5d also shows the strong luminance bias of V3a and MT (Conway and Tsao, 2005).

## Discussion

This study uses a data-driven dimensionality reduction approach, Sparse Supervised Embedding (SSE), to uncover functional signatures for retinotopic visual areas and their organization by eccentricity and upper/lower visual hemifield. The features defining the functional signatures were fMRI responses to gratings varying in hue, saturation, and spatial frequency. The analysis recovered three main components, each representing a multivariate combination of these stimulus features. One component explained the eccentricity representation; the other components separated visual areas and their dorsal/ventral subdivisions. The composition of these components offer insight into the roles of different visual areas, the origin of behavioral asymmetries between upper and lower visual fields, as well as the functional similarity between macaques and humans.

### A functional signature of cortical responses defined by eccentricity

To recover components unrelated to eccentricity, we specified the SSE weight matrix to disregard a voxel’s eccentricity preference. The result was striking: the model discovered two components that discriminated visual areas by their anatomical location (posterior to anterior) and their dorsal/ventral subdivisions (Figure 4). The first component’s weights were predominantly driven by responses to high versus low saturation colors, with more posterior areas showing higher responses to more saturated stimuli. This aligns with previous work in humans indicating that contrast response functions are flatter for more anterior areas (Boynton et al., 1996; Olman et al., 2004; Buracas and Boynton, 2007; Tregillus et al., 2021). The second component discriminated dorsal parcels from ventral parcels of each retinotopic area, primarily based on responses to color, especially daylight-axis colors versus luminance contrast.

Previous work has shown that dorsal V1 has relatively more cortical real estate compared to ventral V1 in monkeys (Van Essen et al., 1984) and humans (Silva et al., 2018; Himmelberg et al., 2022). This asymmetry correlates with higher contrast sensitivity in the lower visual field (Himmelberg et al., 2022), as dorsal V1 maps the ventral visual field. Component 2 here confirms the higher contrast sensitivity among dorsal parcels, as indicated by the highest weighted features distinguishing dorsal from ventral parcels, specifically the three luminance-modulating gratings varying in contrast. But the SSE results cannot be solely explained by higher contrast sensitivity of dorsal parcels, as component 2 also comprises strong weighting of features besides luminance contrast that separate ventral from dorsal parcels. Notably, all features distinguishing ventral from dorsal parcels were related to color modulation, with the two most influential features consisting of daylight-axis colors. We speculate that the dorsal/ventral asymmetry in luminance/color responses reflects a combination of specializations: dorsal parcels for higher contrast sensitivity and ventral parcels for better color detection of the relevant parts of scenes (objects).

Both the SSE and univariate analysis suggest that the primary driver of ventral-dorsal separation is the difference in response between daylight-axis colors and luminance-contrast gratings, rather than simple response differences to color. Asymmetries in color versus luminance responses of dorsal and ventral subdivisions of macaque V4 have been explored by others (Wade et al., 2008). Our results suggest that measuring colors defined by intermediate axes, along with controls for eccentricity, is necessary to uncover the functional signatures that distinguish dorsal versus ventral V4. Dorsal parcels were separated from ventral parcels to a progressively greater extent along the posterior-to-anterior sequence of visual areas. This suggests that the asymmetry in dorsal and ventral V4 responses is inherited and amplified from a dorsal/ventral asymmetry in V1, reminiscent of the amplification of the foveal bias from V1 to V4 (Motter, 2009). We recognize that prior work has not found a dorsal-ventral asymmetry in V1, but one might have been missed because of the low sensitivity of univariate methods compared to multivariate methods, as evident in the present work, where dorsal/ventral asymmetry in V1 did not reach significance in a univariate analysis.

### SSE as a probe of monkey-human cortical homologies

A long-standing question in visual neuroscience is the extent to which macaque and human visual cortical areas are homologous. Homology is well-supported for V1, V2, and V3 (Sereno et al., 1995; DeYoe et al., 1996; Wandell, 1999), and seems supported for inferior temporal areas (Lafer-Sousa et al., 2016). But there are conflicting views on the region in between, centered on V4 (Wade et al., 2002; Fize et al., 2003; Brewer et al., 2005; Larsson and Heeger, 2006; Hansen et al., 2007; Winawer et al., 2010; Roe et al., 2012; Winawer and Witthoft, 2015). The argument for species divergence of V4 has taken two routes. The first concerns the retinotopic organization of the dorsal and ventral candidate regions; the second focuses on the extent to which dorsal versus ventral candidate subdivisions show a bias for color. A cartoon of macaque V4 would show the dorsal division representing the lower visual field and the ventral division representing the upper visual field. This cartoon fails in humans. The candidate ventral region has at least partial representation of the lower visual field (Tootell, 2001; Wade et al., 2002; Brewer et al., 2005; Larsson and Heeger, 2006; Winawer et al., 2010). This failure could be evidence of a species difference. Yet the cartoon also fails for macaques: part of the lower visual field is also represented in ventral V4 (Gattass et al., 1988; Fize et al., 2003). Thus monkeys and humans might not differ in the retinotopic organization of the candidate dorsal and ventral V4 parcels.

What about the second line of reasoning? In humans, candidate dorsal V4 shows relatively lower responsivity to color, while both dorsal and ventral portions of V4 in monkeys supposedly show comparable responses to color (Goddard et al., 2011). But here we find, using the SSE analysis, that dorsal and ventral partitions of monkey V4 diverge in color responsivity, with dorsal regions expected to be less responsive to color than ventral regions. This asymmetry is similar to that in humans. The present results also support the extension to macaques of an observation in humans: ventral partitions across cortical areas generally show relatively greater responses to color (Goddard et al., 2011). The weight of evidence would seem, therefore, to support the idea that V4 is by organization and function similar in humans and macaque monkeys.

### The origin of dorsal / ventral functional asymmetries

What is the origin of the functional asymmetries between dorsal and ventral parcels? One theoretical possibility is that they are inherited from asymmetric retinal photoreceptor distributions. Indeed, the retina shows a subtle asymmetric distribution of cones with polar angle: the inferior retina has slightly higher cone density than the superior retina in both humans (Curcio et al., 1991; Song et al., 2011) and macaques (Perry and Cowey, 1985; Packer et al., 1999), consistent with the higher chromatic responses of the ventral retinotopic parcels. This asymmetry in cone density appears related to rod density. The superior retina has relatively more rods compared to the inferior retina, resulting in a higher relative number of rods to cones (Curcio and Allen, 1990). We wonder if this asymmetry contributes to the contrast response differences observed between dorsal and ventral parcels, given that dorsal parcels show greater responses to luminance contrast, and these parcels receive input from the superior retina with a higher rod-to-cone ratio.

But the organization of the retina cannot fully explain the cortical asymmetries for at least three reasons. First, the retinal ganglion cell density is markedly higher in the superior retina (Curcio and Allen, 1990; Watson, 2014), opposite to the asymmetry in cone density and consistent with the relatively higher acuity of the lower visual field. Second, quantitative work shows that the pattern of cone distribution and other pre-retinal optical factors are grossly insufficient to explain the asymmetries in visual performance (Kupers et al., 2019). Third, differences in cone density do not account for functional asymmetries between dorsal and ventral parcels observed for daylight versus anti-daylight colors. We hypothesize that the dorsal/ventral asymmetries across retinotopic cortex arise from selective sampling of geniculate input by V1, which is then amplified along the cortical hierarchy. Working out the mechanisms underlying these asymmetries will likely benefit from more precise parcellation of the retinotopic representation, including defining parcels by polar angle rather than the coarse partition we used of upper and lower visual field (Himmelberg et al., 2023).

What is the computational objective of the dorsal-ventral asymmetry? It may reflect neural adaptations for processing differences in natural scene statistics between the upper and lower visual field, as in mouse (Denman et al., 2018; Szatko et al., 2020; Qiu et al., 2021). The dorsal-ventral asymmetry we observe in monkeys implies that visual discrimination is more likely to have useful information defined daylight colors in the upper visual field compared to the lower visual field. Daylight-axis colors are important for object recognition and segmentation (Gegenfurtner and Rieger, 2000; Lafer-Sousa et al., 2012; Pearce et al., 2014; Rosenthal et al., 2018), and responses to daylight colors appear to distinguish ventral and dorsal parcels, particularly in V3 and V4 (Figure 5b). A potential behavioral advantage for color-in-service-of-object-vision in the upper visual field may parallel the behavioral advantage in the upper visual field found for face perception (Quek and Finkbeiner, 2014, 2015; Tsurumi et al., 2022) and shape perception (Zito et al., 2016) in the upper visual field.

### Broader applications of the SSE method

Neural tuning properties differentiate retinotopic areas, as discovered using parametrically generated texture stimuli (Freeman et al., 2013; Ziemba et al., 2016). Building on the conceptual framework established by Hubel and Wiesel, which considers complex stimulus selectivity as a chain of computations starting with punctate center-surround receptive fields of retinal ganglion cells, our work uses SSE to determine whether visual areas, not just neurons, can be characterized by functional signatures constructed from simple building blocks. The present results serve as a proof-of-principle, indicating that SSE could be fruitfully applied to identify functional signatures using different building blocks beyond gratings. For instance, we aim to use this approach to identify abstract rules common to various subregions (color-biased domains, face patches, place-biased domains) within each of the four stages of inferior temporal cortex.

## Acknowledgments

This work was supported by the Intramural Research Program of the National Eye Institute. We thank L. Wald and A. Mareyam for providing the four-channel magnetic resonance coil. We also thank R. Lafer-Sousa for help with MRI scanning. The original MRI scanning was carried out at the Athinoula A. Martinos Center for Biomedical Imaging at the Massachusetts General Hospital, using resources provided by the Center for Functional Neuroimaging Technologies and a P41 Biotechnology Resource grant supported by the National Institute of Biomedical Imaging and Bioengineering (P41EB015896). This work also involved the use of instrumentation supported by the US National Institutes of Health Shared Instrumentation Grant Program and/or High-End Instrumentation Grant Program (S10RR021110).

